# Design and immunogenicity of an HIV-1 clade C pediatric envelope glycoprotein stabilized by multiple platforms

**DOI:** 10.1101/2024.09.14.613016

**Authors:** Sanjeev Kumar, Iván del Moral-Sánchez, Swarandeep Singh, Maddy L. Newby, Joel D. Allen, Tom P. L. Bijl, Yog Vaghani, Liang Jing, Eric A. Ortlund, Max Crispin, Anamika Patel, Rogier W. Sanders, Kalpana Luthra

## Abstract

Various design platforms are available to stabilize soluble HIV-1 envelope (Env) trimers, which can be used as antigenic baits and vaccine antigens. However, stabilizing HIV-1 clade C trimers can be challenging. Here, we stabilized an HIV-1 clade C trimer based on an Env isolated from a pediatric elite-neutralizer (AIIMS_329) using multiple platforms, including SOSIP.v8.2, ferritin nanoparticles (NP) and an I53-50 two-component NP, followed by characterization of their biophysical, antigenic, and immunogenic properties. The stabilized 329 Envs showed binding affinity to trimer-specific HIV-1 broadly neutralizing antibodies (bnAbs), with negligible binding to non-neutralizing antibodies (non-nAbs). Negative-stain electron microscopy (nsEM) confirmed the native-like conformation of the Envs. Multimerization of 329 SOSIP.v8.2 on ferritin and two-component I53-50 NPs improved the overall affinity to HIV-1 bnAbs and immunogenicity in rabbits. These stabilized HIV-1 clade C 329 Envs demonstrate the potential to be used as antigenic baits and as components of multivalent vaccine candidates in future.

## INTRODUCTION

Clade C accounts for approximately 50% of HIV-1 infections worldwide and is responsible for more than 90% of infections in India and South Africa ^1,2^. HIV-1 envelopes (Env) isolated from elite-neutralizer individuals who develop broadly neutralizing antibodies (bnAbs) can inform HIV-1 vaccine design by serving as templates for the induction of similar bnAb responses through vaccination ^3–8^. The design and development of stabilized native-like HIV-1 Env soluble trimer antigens, predominantly from non-clade C isolates, have enabled the induction of neutralizing antibody (nAb) responses against HIV-1 in animal models ^9–13^. Furthermore, native-like Envs have been used as antigenic baits to identify exceptionally potent second-generation HIV-1 bnAbs from both adults and children ^5,14–17^. However, the instability and low expression levels of HIV-1 clade C Env trimers in soluble form have hindered the development of clade C Env based HIV-1 vaccines to induce protective bnAb responses ^18–20^. While it seems unlikely that single Env-based regimens will suffice to induce bnAb responses, sequential immunizations with multivalent immunogens or cocktails of different Envs hold greater potential ^8,21^. This underlies the need to generate stable Env trimers from HIV-1 strains of diverse geographical origins and distinct clades, particularly those isolated from elite-neutralizers who develop exceptionally potent bnAbs with multi-epitope specificities ^4,20,22,23^.

The SOSIP mutations are well-known to produce native-like soluble HIV-1 Env trimers ^9,10,10,24,25^. However, the use of these mutations for the design of clade C Envs has mostly yielded low expressing Envs with poor antigenicity and immunogenicity ^19,20^. In the past few years, various studies described mutations that increase the purification yield, antigenicity and stability of soluble Env proteins, including some clade C Envs ^11,12,26–31^. Most of these novel mutations have been defined using the clade A BG505 Env ^10,29–31^. However, the effects of these mutations on the other clade specific Envs is often limited. Moreover, a limited number of HIV-1 clade C native-like Envs from India have been described thus far ^19,20^ and none from pediatric elite-neutralizers ^5,22,23,32^.

We previously reported the characterization of Env sequences obtained from a pair of Indian clade C chronically infected pediatric elite-neutralizer monozygotic twins (AIIMS_329 and AIIMS_330), whose plasma exhibited exceptionally strong bnAb responses with multiple epitope specificities against a large panel of multi-clade heterologous Env pseudoviruses ^5,22^. Such Envs have the potential as templates for stabilization and for immunogen design, and they could serve as useful antigenic baits to isolate HIV-1 bnAbs for immunotherapeutic purposes. This is reinforced by the fact that the most studied HIV-1 Env sequence, BG505, was isolated from an infant transmitted founder virus and multiple BG505-derived native-like Env trimers are currently being evaluated in vaccine clinical trials ^8,13,21,24,28^. In the past few years, using BG505 trimers as an antigenic bait, several second-generation potent HIV-1 bnAbs have been identified from both adults and children ^5,14–16,33^.

Herein, we designed and characterized a soluble HIV-1 clade C_329 Env trimer, derived from a circulating virus in an Indian pediatric elite neutralizer AIIMS_329, by stabilizing the sequence using SOSIP v8.2 ^24^, displaying it on ferritin ^34^ and self-assembling two-component I53-50 nanoparticles (NPs) ^27,35^. The stabilized native-like 329 SOSIP.v8.2 Env trimer showed high binding to most HIV-1 bnAbs and negligible binding to non-neutralizing antibodies (non-nAbs). Multimerization of 329 SOSIP.v8.2 trimers on ferritin and two-component I53-50 NPs improved the affinity to HIV-1 bnAbs and immunogenicity in rabbits. Native-like conformation of the proteins was confirmed by low-resolution negative-stain electron microscopy (nsEM) and cryo-electron microscopy (cryoEM). We have for the first time, stabilized and demonstrated the immunogenicity of varied versions of an Indian Clade C native-like Env trimer, derived from a pediatric elite neutralizer; with a potential to be used as a template for a clade C based vaccine and as a bait to isolate HIV-1 bnAbs for therapeutic / prophylactic purposes. Multimeric antigen presentation has evolved as a promising strategy, that can be applied in the future to improve the overall stability and antigenicity of other (unstable) HIV-1 clade C and non-clade C trimeric Env glycoproteins.

## RESULTS

### Design and biophysical characterization of a native-like 329 SOSIP.v8.2 Env trimer

We previously reported the isolation and characterization of multiple Env pseudoviruses from an Indian HIV-1 clade C seropositive pediatric elite-neutralizer (AIIMS_329), whose plasma antibodies showed broad and potent HIV-1 neutralization in a longitudinal study ^22^. One of these autologous Env pseudoviruses, 329.14.B1, showed exceptional susceptibility to neutralization by the majority of bnAbs in a panel covering multiple epitope specificities, and was resistant to non-nAbs and sCD4 ^22^. Hence, we selected this Env to stabilize it in soluble trimeric form. We engineered the 329 SOSIP.v8.2 Env trimer (**Fig. 1A**) by introducing SOSIP mutations (501C-605C, 559P), including a multibasic furin cleavage site (hexa-arginine or R6) between gp120 and gp41. This protein also incorporated TD8 (47D, 49E, 65K, 165L, 429R, 432Q, 500R) ^11^ and MD39 stabilizing mutations (106E, 271I, 288L, 304V, 319Y, 363Q, 519S, 568D, 570H, 585H) ^31^, as well as a mutation to reduce V3-exposure (66R) ^25^. We also introduced changes to optimize the epitope of PGT145 (166R, 168K, 170Q, 171K) (**Fig. S1)**.

**Figure 1:**
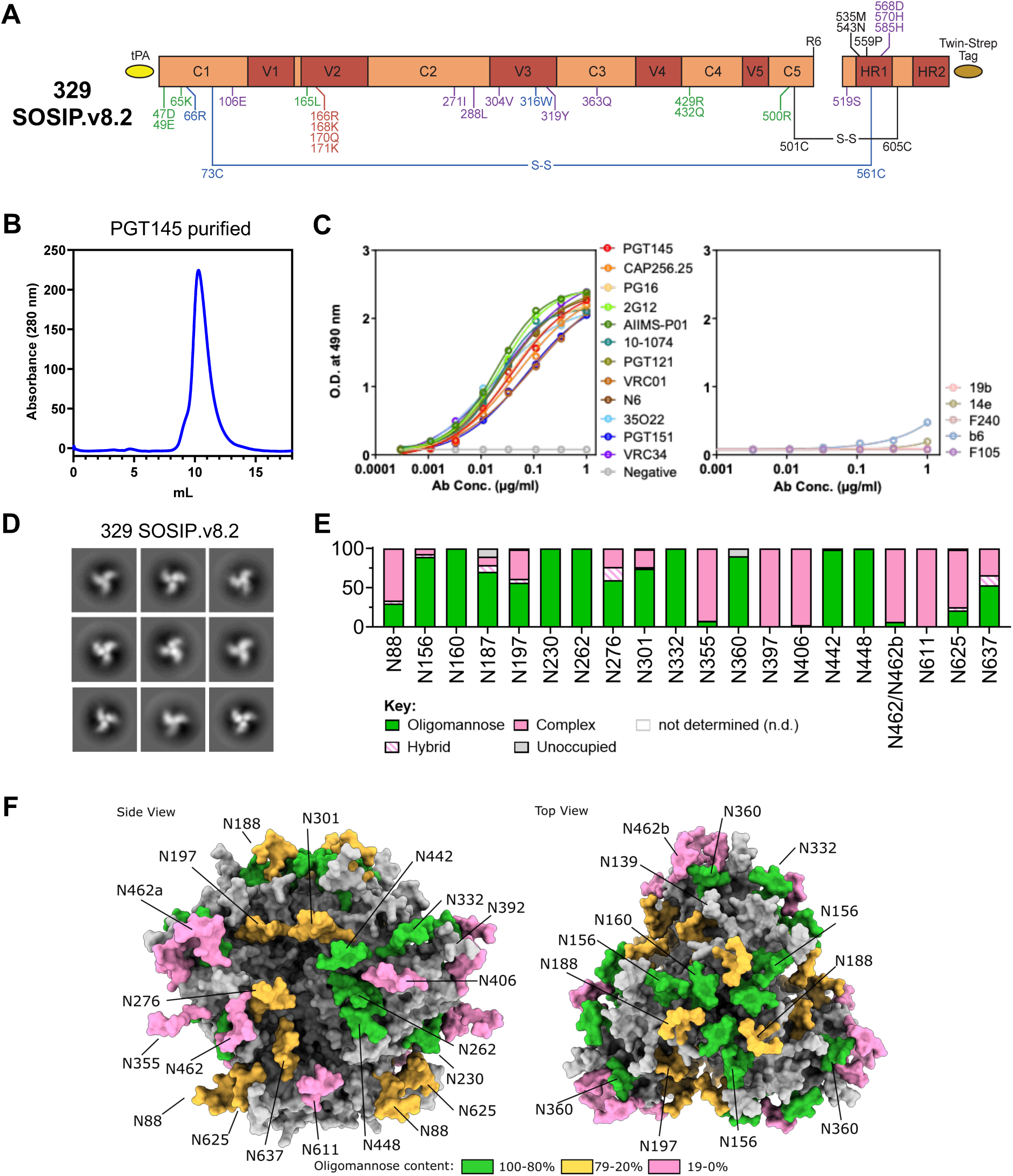
Design and biophysical characterization of the clade C 329 SOSIP.v8.2 Env trimer. **A.** Linear representation of the 329 SOSIP.v8.2 construct, with SOSIP.664 mutations (501C-605C, 559P, R6) in black, further stabilizing SOSIP mutations (66R, 316W, 73C-561C) in blue, TD8 mutations (47D, 49E, 65K, 165L, 429R, 432Q, 500R) in green, MD39 mutations (304V, 319Y, 363Q, 519S, 568D, 570H, 585H) in purple, and PGT145 epitope modifications (166R, 168K, 170Q, 171K) in red. **B.** SEC profile of PGT145-purified 329 SOSIP.v8.2 on a Superdex 200 Increase 10/300 GL column. **C.** StrepTactinXT ELISA assay with PGT145-purified 329 SOSIP.v8.2 against a panel of bNAbs (left) and non-NAbs (right). **D.** 2D class averages generated from nsEM data of the PGT145-purified 329 SOSIP.v8.2 protein. **E.** Site-specific glycan analysis of PGT145-purified 329 SOSIP.v8.2 protein. Values represented are specified in Table S1. PNGS are displayed as aligned with HxB2. Data could not be determined (n.d.) for sites N143, N241 and N397. The glycan modifications on the remaining sites were classified into three categories: high mannose (corresponding to any composition containing two HexNAc residues, or three HexNAc and at least 5 hexoses), complex, or unoccupied. The proportion of peptides and glycopeptides corresponding to each of these categories was colored green for high mannose, pink for complex and grey for unoccupied. **F.** Model of the glycan shield of 329 SOSIP.v8.2 generated using AlphaFold 3 and Re-Glyco. A representative Man_5_GlcNAc_2_ glycan is modelled at each site and colored according to the % oligomannose-type glycans displayed in panel E. Sites that could not be determined are colored grey.

We expressed 329 SOSIP.v8.2 in HEK293F cells, followed by PGT145 antibody affinity chromatography purification (**Fig. 1B**). The purification yield of this trimer was ∼0.6 mg/L which is comparable (0.4 – 0.6 mg/ml) to a previous clade C Env from South African strain (CZA97.012, 0.4 – 0.6 mg/mL)^36^. The 329 SOSIP.v8.2 Env showed a single gp140 Env trimer band in BN-PAGE (**Fig S2A**). In ELISA binding assays, 329 SOSIP.v8.2 Env interacted well with HIV-1 bnAbs and showed negligible binding to all non-nAbs tested, consistent with a native-like closed conformation (**Fig 1C**). The native-like conformation of stabilized 329 SOSIP.v8.2 Env well-ordered trimers was further confirmed by nsEM (**Fig 1D**). To determine the glycan composition of 329 SOSIP.v8.2 Env trimer, we performed site-specific glycan analysis by mass spectrometry. Overall, the glycosylation profile of the 329 SOSIP.v8.2 clade C Env trimer presents a similar abundance of oligomannose-type glycan signatures at canonical “mannose sites”, including N160, N262, N332 and N448, as compared to previously characterized clade A and B Envs ^26–28^, (**Fig. 1E, S3 and Table S1**). To display the glycan holes or absence of PNGS sites (N289, N295, N339 and N386) in the 329 Env, a 3D model of the 329 SOSIP.v8.2 Env representing glycosylation was generated using AlphaFold 3^37^, GlycoShape and Re-Glyco^38^ (**Fig. 1F**). Overall, the biophysical characterization suggests that 329 SOSIP.v8.2 Env is successfully stabilized in a soluble native-like state and efficiently displays the epitopes for all known HIV-1 bnAbs tested in this study.

### Displaying 329 SOSIP.v8.2 Env on ferritin and two-component nanoparticles

To increase the valency of 329 Env, we presented 329 SOSIP.v8.2 trimers on the surface of protein nanoparticles (NPs). First, we fused the C-terminus of 329 SOSIP.v8.2, after position 664, to the N-terminus of a previously described *H. pylori* ferritin ^34^ (GenBank accession no. NP_223316), starting at position Asp5, using a flexible Gly-Ser linker (GSG) (**Fig. 2A**). The ferritin NPs can display 8 native-like Env trimers ^34^. Second, we previously described the computational design of two-component self-assembling I53-50 NPs ^35^, which are 120-subunit assemblies of icosahedral symmetry comprising 20 trimeric (I53-50A) and 12 pentameric (I53-50B) subunits. Therefore, each I53-50 NP can present up to 20 trimeric antigens fused to the I53-50A components. The possibility to purify the I53-50A-antigen fusion proteins with trimer-selective purification methods before in vitro assembly with the I53-50B component ensures the presentation of native-like trimers exclusively. To further increase the valency of the 329 SOSIP.v8.2 Env, we genetically fused it to I53-50A component (I53-50A.1NT1) via a Gly-Ser-rich linker (GSGGSGGSGGSGGS) (**Fig. 2B**).

**Figure 2:**
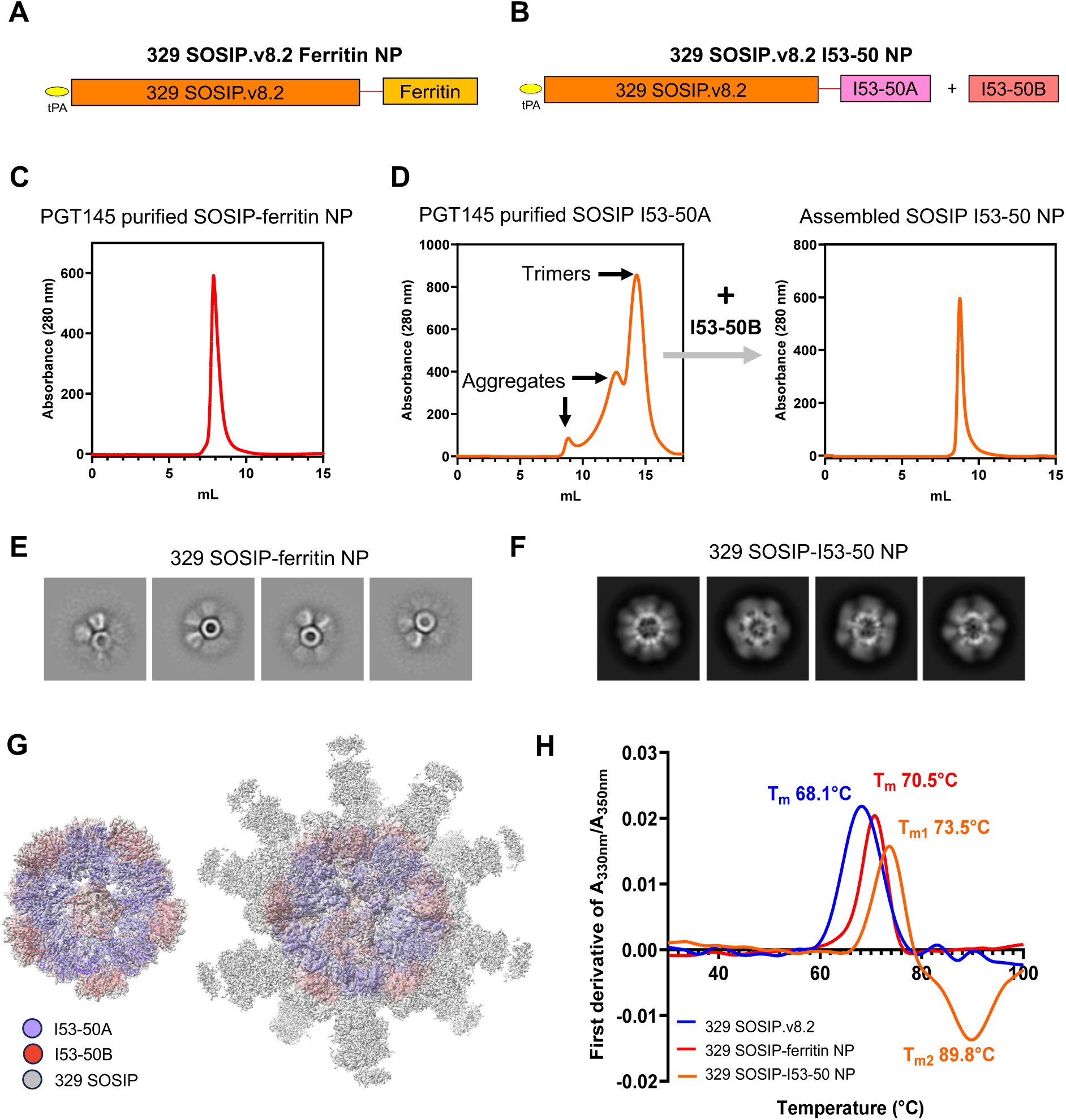
Design and biophysical characterization of 329 SOSIP-ferritin and SOSIP-I53-50 nanoparticles. **A, B.** Linear representations of the 329 SOSIP-ferritin (A) and 329 SOSIP-I53-50A (B) fusion proteins. The red lines represent GSG and GSGGSGGSGGSGGS flexible linkers. **C.** SEC profile of PGT145-purified 329 SOSIP-ferritin NPs on a Superdex 200 Increase 10/300 GL column. **D.** SEC profile of PGT145-purified 329 SOSIP-I53-50A fusion protein (left) and 329 SOSIP_I53-50 assembled NPs (right) on a Superose 6 Increase 10/300 GL column. **E, F.** nsEM-generated 2D class averages of 329 SOSIP-ferritin (**E**) and 329 SOSIP-I53-50 (**F**) NPs. **G.** 3.8 Å resolution cryo-EM map showing details of I53-50 NP core (left) and density for 329 SOSIP trimers displayed on I53-50 NP core (right) can be seen at low contour level. **H.** Denaturing profiles of 329 SOSIP, 329 SOSIP-ferritin NPs and 329-SOSIP-I53-50 NPs, obtained by nanoDSF and used to determine the T_m_ values referred to in the results section.

Next, the resulting SOSIP-ferritin and SOSIP-I53-50A fusion proteins were expressed in HEK293F cells, followed by purification using PGT145 bnAb-affinity chromatography (**Fig. 2C and 2D**). The purified 329 SOSIP-I53-50A was mixed with I53-50B component, to fully assemble the 329 SOSIP-I53-50 NPs (**Fig. 2D**). SEC purification of the fully assembled 329 SOSIP-I53-50 NP revealed a peak at an elution volume of ∼9.0 which is further shifted compared to 329 SOSIP-I53-50A peak, indicating formation of high-molecular-weight complex (**Fig. 2D, right panel**). The purified 329 SOSIP-ferritin and SOSIP-I53-50 NPs showed a single gp140 Env-displaying NPs band in BN-PAGE (**Fig S2B and S2C**). The purification yields for SOSIP-ferritin and assembled SOSIP-I53-50 NPs were 0.6 mg/mL and ∼1.0 mg/L, respectively. Fully assembled NPs when imaged by nsEM show well assembled NP structures with 329 SOSIP.v8.2 Env trimer attached to the NP core (**Fig. 2E and 2F**). These results confirmed the multimeric presentation of 329 SOSIP.v8.2 Env trimer on self-assembled ferritin and two-component I53-50 NPs. To confirm the proper assembly of I53-50 NP and their structural integrity, we performed single-particle cryoEM analysis on 329 SOSIP-I53-50 NPs. The 3D model generated from the cryoEM data confirmed the construction of I53-50 core (left) and multimeric presentation of 329 SOSIP trimers attached to each I53-50A moiety (right) (**Fig. 2G**). Due to high flexibility in the linker between the two domains, the 329 SOSIP trimers were poorly resolved and appear as diffused densities surrounding the well-resolved I53-50NP core of ∼3.8 Å resolution.

### 329 Env NPs show improved thermostability and antigenicity

Next, we were interested to determine the effect of multimeric display of 329 SOSIP.v8.2 on the stability and antigenic properties of self-assembled 329 SOSIP-ferritin and 329 SOSIP-I53-50 NPs. First, we evaluated the thermostability of the proteins by nano differential scanning fluorimetry (nanoDSF). Both SOSIP-ferritin and SOSIP-I53-50 NPs presented a higher T_m_ of 70.5°C and 73.5°C, respectively, as compared to the T_m_ of 68.1°C observed for 329 SOSIP.v8.2 Env trimer (**Fig. 2H**). Next, we determined their antigenicity using Bio-Layer Interferometry (BLI). The V2-apex-targeting bnAbs PGT145 and CAP256.25 showed enhanced binding to both 329 SOSIP-I53-50 and SOSIP-ferritin NPs compared to soluble 329 SOSIP.v8.2 Env trimer (**Fig. 3**). HIV-1 gp120-gp41 interface targeting bnAb PGT151 showed slightly lower binding to assembled NPs, consistent with the lower accessibility of base-proximate epitopes of SOSIP trimers multimerized on NPs ^26,27^. Furthermore, the CD4bs-specific and N332-supersite-specific bnAbs VRC01 and PGT121 interacted more efficiently with 329 SOSIP-I53-50 NPs, compared to the soluble and ferritin-displayed counterparts. None of the non-nAbs tested interacted with the soluble and multimerized 329 SOSIP.v8.2 trimers, including b6 and F105 against the CD4bs, and 19b targeting the V3 (**Fig. 3**). Overall, these findings suggest that 329 SOSIP.v8.2 trimers maintain their native-like conformation and improve their overall antigenicity when multimerized on ferritin and I53-50 NPs.

**Figure 3:**
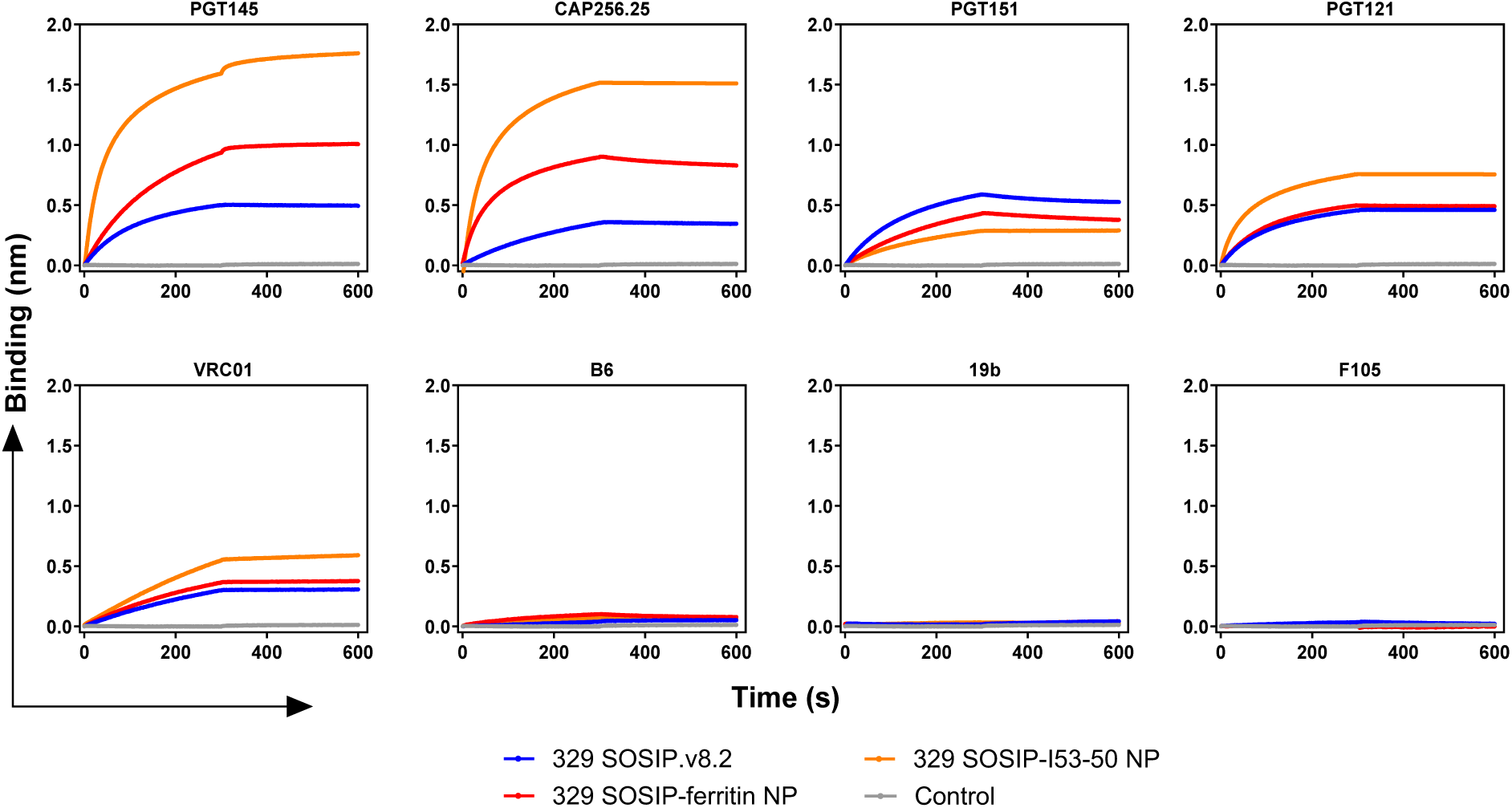
Antigenic analysis of 329 SOSIP, SOSIP-ferritin and SOSIP-I53-50 nanoparticles. ProtA BLI assay with 329 SOSIP, 329 SOSIP-ferritin NPs and 329-SOSIP-I53-50 NPs and a panel of bnAbs (PGT145, CAP256.25, PGT151, PGT121, VRC01) and non-nAbs (B6, 19b, F105). The experiment was performed in duplicate, and the curves shown correspond to one of these repetitions.

**Figure 4:**
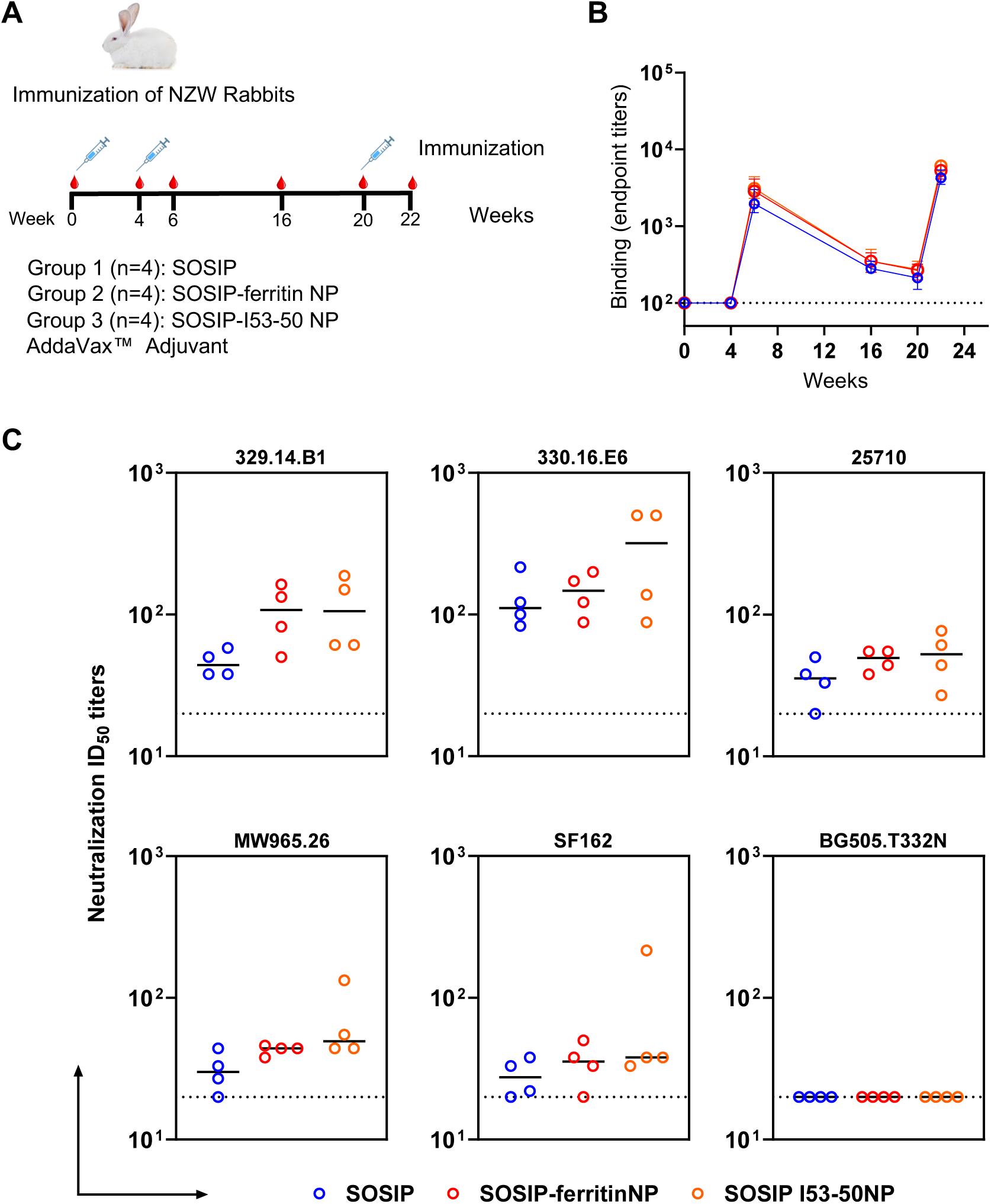
Immunogenicity of 329 SOSIP.v8.2 Env trimer, SOSIP-ferritin and SOSIP-I53-50 NPs in rabbits. **A.** Rabbit immunization schedule. Four groups of New Zealand White rabbits were immunized at weeks 0, 4, and 20 with 10 µg of SOSIP trimer (Group 1, n=4) or equimolar amounts of SOSIP-ferritin NPs (25 µg) (Group 2, n=4) or SOSIP-I53-50 NPs (32 µg) (Group 3, n=4) and placebo (PBS with Adjuvant) (Group 4, n=2). Antibody responses were evaluated at weeks 0, 4, 6, 16, 20, and 22. **B.** Endpoint antibody binding titers over time against 329 SOSIP.v8.2 trimer as measured by StreptactinXT ELISA. Dots and error bars represent the median binding titers and standard deviations. No significant differences were found by a Kruskal–Wallis statistical test between groups at any timepoint tested. **C.** Midpoint neutralization titers (ID_50_) for week 22 sera of the immunized rabbits against a panel of pseudoviruses. Horizontal lines represent the geometric means of ID_50_ titers. No significant differences between groups were found by an unpaired two-tailed Mann–Whitney U-test. Assay cut-off is marked with a dotted line.

### Multimerization on NPs improves the immunogenicity of 329 Env trimers in rabbits

Next, we compared the immunogenicity of 329 SOSIP.v8.2 trimers with its 329 SOSIP-ferritin and 329 SOSIP-I53-50 NPs variants in New Zealand White (NZW) rabbits. Four groups of four female rabbits were immunized at weeks 0, 4, and 20 with 20 µg of 329 SOSIP, or the equimolar amount presented on 329 SOSIP-ferritin and 329 SOSIP-I53-50 NPs, formulated in AddaVax™ adjuvant. Sera were collected from the rabbits at weeks 0, 4, 6, 16, 20 and 22 to assess the antibody responses (**Fig. 3A**). First, we measured the binding titers of the collected immune sera at the different timepoints to a 329 SOSIP.v8.2 trimer in ELISA (**Fig. 3B**). An increase in binding titers occurred at week 6 and week 22, two weeks after the first and second booster immunizations, respectively (**Fig. 3B**). We detected no significant differences in the binding titers induced by the different immunogens at any timepoint. Moreover, we evaluated the development of neutralizing antibody (nAb) titers against autologous AIIMS_329 (329.14.B1), closely related AIIMS_330 (330.16.E6) and heterologous clade C (25710 and MW965.26), Tier 1 clade B (SF162), and Tier 2 clade A (BG505.T332N) pseudoviruses (**Fig. 3C and Table S2**). Presentation on both ferritin and I53-50 NPs resulted in improved nAb titers against the autologous 329.14.B1 and the closely related 330.16.E6 pseudoviruses, compared to the soluble format, although the differences were not statistically significant. Neutralizing responses to the Tier 1A MW965.26 virus are dominated by V3-specific nAbs, unable to neutralize most Tier 2 primary HIV-1 isolates. The three 329 immunogens tested induced similarly low MW965.26-neutralizing responses, indicating effective masking of V3. None of the 329 immunogens induced BG505-neutralizing responses, consistent with the low BG505 nAb activity observed in the plasma of the AIIMS_329 HIV-1 infected elite neutralizer ^22^. Overall, these results demonstrate that both soluble and NP-displayed 329 SOSIP trimers effectively induce autologous nAb responses and that presenting them on NPs improves the nAb responses induced.

## DISCUSSION

In general, clade C HIV-1 Env trimers are inherently unstable and difficult to stabilize for vaccine or capture reagent use ^11,12,18,20^. A majority of the native-like soluble HIV-1 clade C trimers have been stabilized using cleavage-independent single-chain or native-flexibly linked (NFL) modifications ^11,12^. However, these platforms have shown decreased binding to gp120-41 interface targeting bnAbs (e.g. PGT151, ACS202, VRC34), making them less suitable for use as an antigenic-bait to discover novel bnAbs. HIV-1 Envs from elite-neutralizers can be critical templates to design vaccine candidates ^3,4,20^. Discovering novel HIV-1 bnAbs from clade C infected donors and evaluating the quality, breadth and epitope specificity of the bnAbs induced by geographically distinct HIV-1 Env sequences obtained from elite-neutralizers, including children, could provide critical insights for effective HIV-1 vaccine design and vaccination strategies. We previously identified and characterized HIV-1 clade C Envs from a pair of chronically infected monozygotic twin pediatric elite-neutralizers, AIIMS_329 and AIIMS_330. An HIV-1 Env pseudovirus from AIIMS_329 showed susceptibility to majority of the HIV-1 bnAbs and resistance to non-nAbs and sCD4 ^22^. In the past recent years, multimeric presentation of class I fusion viral Env glycoproteins e.g. Respiratory Syncytial Virus (RSV) F protein, influenza hemagglutinin (HA), Lassa virus (LASV) GPC protein, SARS-CoV-2 S protein and HIV-1 Env, on ferritin and two-component protein nanoparticles (NPs) has enabled efficient controlled production of well-ordered NP-based vaccine candidates ^26–28,34,39–41^.

Herein, we describe the design of stabilized native-like 329 Env trimers using multiple vaccine platforms, including soluble, ferritin and two-component NPs. We introduced newly reported HIV-1 Env stabilizing mutations that are known to increase the purification yield, antigenicity and stability of soluble Env proteins ^11,12,26–31^. The 329 SOSIP.v8.2 Env, SOSIP-ferritin and SOSIP-I53-50 two-component NPs were efficiently expressed and purified using PGT145 bnAb-affinity chromatography. The purified Env and NPs showed reactivity to HIV-1 bnAbs and negligible binding to non-nAbs. The 329 SOSIP.v8.2 Env showed closed native-like conformation in nsEM analysis. Immunization studies in rabbits demonstrated the improved immunogenicity by both ferritin and two-component NPs as compared to 329 SOSIP.v8.2 Env.

Although a large number of HIV-1 Env immunogens have been stabilized and characterized using multiple platforms using cleavage-independent (single-chain, NFL, and UFO) or cleavage dependent (SOSIP) versions from distinct viruses, the majority of the studies were based on a clade A based transmitted-founder Env isolated from a 6-week old infant BG505 ^10–13,24,25,29,30,42^. To the best of our knowledge, no such Envs were designed and characterized from an HIV-1 chronically infected pediatric elite-neutralizer in native-like soluble form.

In our immunization studies, we observed relatively low neutralization ID_50_ titers, plausibly due to reduced antigenic exposure, the presence of more complex glycan composition in 329 Env or because of the use of the AddaVax adjuvant. Similar to the results in a recent study on highly thermostable BG505 Env trimers, we observed weak neutralization of MW965.26 clade C Tier 1 virus in the animals vaccinated with 329 SOSIP Env or its NP-fusion proteins ^30^. This could be an effect of non-native mutations in the 329 Env that were introduced according to our stabilizing strategies, including suppression of the immunodominant V3 and CD4i epitopes. Further, we detected lesser neutralizing activity of the immune sera of rabbits, immunized with 329 SOSIP-I53-50 NPs, for the autologous AIIMS_329 HIV-1 Env pseudovirus in comparison to a closely related AIIMS_330 Env pseudovirus, which could plausibly be due to the better exposure of neutralizing determinants on the AIIMS_330 virus ^5,22^. Detailed studies to understand the effect of different adjuvants, then role of complex glycans and modulating various Env mutations are warranted to study their subsequent effects on immunogenicity.

In conclusion, our findings demonstrate that multimeric presentation of HIV-1 clade C trimer could improve its overall stability, antigenicity, and immunogenicity, through display of homogeneous arrays of native-like HIV-1 Env trimers. As, HIV-1 clade C accounts for fifty percent of the HIV infections worldwide, it is critical to discover and characterize novel bnAbs from both adults and children to determine their ability to effectively neutralize HIV-1 clade C circulating in the infected individuals of varied age group in a population and confer protection. Further, design and characterization of geographically distinct clade C Env with ability to elicit protective elite HIV-1 neutralizing bnAb responses during the course of natural infection are critical to further guide the rational design and development of globally effective vaccine candidates. The 329 SOSIP.v8.2 Env designed and stabilized in this study could serve as a suitable antigenic-bait. The thermostable 329 SOSIP-ferritin and 329 SOSIP-I53-50 NPs can be components of multivalent immunogens aimed to elicit multiclade specific or broad neutralizing antibody responses to HIV-1 in future, especially in the population of low- or middle-income countries and can facilitate the vaccine distribution without any requirement of maintaining the cold-chain storage conditions.

## METHODS

### Construct design

The 329 *env* gene was derived from a previously identified Indian clade C HIV-1 sequence obtained from a pediatric elite-neutralizer AIIMS_329 (329.14.B1, GenBank: MK076593.1), as previously described^22^.

The 329 SOSIP.v8.2 construct was designed by incorporating the SOSIP mutations (501C-605C, 559P), including a multibasic furin cleavage site (hexa-arginine or R6) between gp120 and gp41. This protein also incorporated TD8 (47D, 49E, 65K, 165L, 429R, 432Q, 500R) and MD39 stabilizing mutations (106E, 271I, 288L, 304V, 319Y, 363Q, 519S, 568D, 570H, 585H), as well as a mutation to reduce the V3-exposure (66R)^21^. We also introduced changes to optimize the epitope of PGT145 (166R, 168K, 170Q, 171K) (**Fig. S1**).

The 329.SOSIP.v8.2-ferritin construct was generated by fusing the N-terminus from *Helicobacter pylori* ferritin (Genbank accession no. NP_223316), starting from Asp5, to the SOSIP.664 C-terminus (truncated at position 664), separated by a Gly-Ser (<u>GS</u>G) linker, as described previously^28^.

To create the 329-I53-50A.1NT1 construct, the original I53-50A.1NT1 plasmid was described previously^26,27^. Modifications constitute the introduction of <u>GS</u>LEHHHHHH after the final residue to introduce a C-terminal histidine-tag.

All constructs comprised the above-described sequences preceded by a tissue plasminogen activator (tPA) signal peptide (MDAMKRGLCCVLLLCGAVFVSPSQEIHARFRRGAR). Untagged Env constructs presented a STOP codon after position 664. Strep-tagged SOSIP.v8.2 constructs included an additional Twin-Strep-Tag amino acid sequence (<u>GS</u>GGSSAWSHPQFEKGGGSGGGSGGSAWSHPQFEKG) after position 664. In every case, the underlined GS residues were encoded by a BamHI restriction site useful for cloning purposes.

All genes were codon-optimized for mammalian expression and synthesized by Genscript (Piscataway, USA), and cloned by restriction-ligation into a pPI4 plasmid.

### HIV-1 envelope protein expression

SOSIP Env and SOSIP Env-NP fusion proteins were expressed as described previously^29,30^. Briefly, HIV-1 Env and furin protease-encoding plasmids were mixed in a 3:1 Env to furin ratio (w/w) and incubated with PEImax (Polysciences Europe GmBH, Eppelheim, Germany) in a 3:1 (w/w) PEImax to DNA ratio. Subsequently, the transfection mixtures were added to the supernatant of HEK293F suspension cells (Invitrogen, cat no. R79009), maintained in FreeStyle Expression Medium (Gibco) at a density of 0.8–1.2 million cells/mL. Seven days post-transfection, supernatants were harvested, centrifuged, and filtered using Steritops (0.22 µm pore size; Millipore, Amsterdam, The Netherlands) before protein purification.

### HIV-1 envelope protein purification

SOSIP Env and SOSIP Env-NP fusion proteins were purified by PGT145 immunoaffinity chromatography as described earlier^29,30^. Briefly, unpurified proteins contained in HEK293F filtered supernatants were captured on PGT145-functionalized CNBr-activated sepharose 4B beads (GE Healthcare) by overnight rolling incubation at 4 °C. Subsequently, the mixes of supernatant and beads were passed over Econo-Column chromatography columns (Biorad). The columns were then washed with three column volumes of a 0.5 M NaCl and 20 mM Tris HCl pH 8.0 solution. After elution with 3 M MgCl_2_ pH 7.5, proteins were buffer exchanged into TN75 (75 mM NaCl, 20 mM Tris HCl pH 8.0) or PBS buffers by ultrafiltration with Vivaspin20 filters (Sartorius, Gӧttingen, Germany) of MWCO 100 kDa. Protein concentrations were determined from the A280 values measured on a NanoDrop2000 device (Thermo Fisher Scientific) and the molecular weight and extinction coefficient values calculated by the ProtParam Expasy webtool.

### I53-50B.4PT1 protein expression and purification

I53-50B.4PT1 protein purification was performed as described earlier^26,27^. Briefly, Lemo21 cells (DE3) (NEB), which were grown in LB (10 g Tryptone, 5 g Yeast Extract, 10 g NaCl) in 2 L baffled shake flasks or a 10 L BioFlo 320 Fermenter (Eppendorf), were transformed with a I53-50B.4PT1-encoding plasmid. After inducing protein expression by the addition of 1 mM IPTG, cells were subjected to shaking for ∼16 h at 18 °C. Microfluidization was used to harvest and lyse the cells, using a Microfluidics M110P machine at 18,000 psi in 50 mM Tris, 500 mM NaCl, 30 mM imidazole, 1 mM PMSF, 0.75% CHAPS. Proteins were purified by applying clarified lysates to a 2.6×10 cm Ni Sepharose 6 FF column (Cytiva) on an AKTA Avant150 FPLC system (Cytiva). A linear gradient of 30 mM to 500 mM imidazole in 50 mM Tris, pH 8, 500 mM NaCl, 0.75% CHAPS was used to elute both proteins. Next, the pooled fractions were subjected to size-exclusion chromatography on a Superdex 200 Increase 10/300, or HiLoad S200 pg GL SEC column (Cytiva) in 50 mM Tris pH 8, 500 mM NaCl, 0.75% CHAPS buffer. I53-50B.4PT1 elutes at ∼0.45 CV. Prior to nanoparticle assembly, protein preparations were tested to confirm low levels of endotoxin. To remove endotoxin, purified I53-50B.4PT1 was immobilized on Ni^2+^-NTA resin in a 5 mL HisTrap HP column (GE Healthcare) equilibrated with the following buffer: 25 mM Tris pH 8, 500 mM NaCl, 0.75% CHAPS. Immobilized I53-50B.4PT1 was then washed with ∼10 CV of the equilibration buffer. The protein was eluted over gradient to 500 mM imidazole in equilibration buffer. Fractions containing I53-50B.4PT1, which elutes around ∼175 mM imidazole, were concentrated in a Vivaspin filter with a 10 kDa molecular weight cutoff and subsequently dialyzed twice against equilibration buffer (GE Healthcare).

### HIV-1 SOSIP-I53-50NP assembly

HIV-1 SOSIP-I53-50NP assembly was performed as described earlier^26,27^. Briefly, after PGT145-purification (see HIV-1 Env protein expression and purification), the SOSIP-component A fusion protein (329 SOSIP.v8.2-I53-50A.1NT1) was passed through a Superose 6 Increase 10/300 GL (GE Healthcare) SEC column in Assembly Buffer II (25 mM Tris, 500 mM NaCl, 5% glycerol pH 8.2) to remove aggregated proteins. The glycerol component was included in the Assembly Buffer II to minimize aggregation of the SOSIP-component A fusion proteins during the assembly of NPs, but we found that their presence increased the recovery of the assembled NPs during the concentration and dialysis stages described below. After the SEC procedure, the column fractions containing non-aggregated SOSIP-I53-50A.1NT1 proteins were immediately pooled and mixed in an equimolar ratio with I53-50B.4PT1 (produced as described above) for an overnight (∼16 h) incubation at 4 °C. The assembly mix was then concentrated at 350 × *g* using Vivaspin filters with a 10 kDa molecular weight cutoff and passed through a Superose 6 Increase 10/300 GL column in Assembly Buffer II (GE Healthcare). The fractions corresponding to the assembled NPs (elution between 8.5 and 10.5 mL with a peak at 9 mL) were pooled and concentrated at 350 × *g* using Vivaspin filters with a 10 kDa molecular weight cutoff (GE Healthcare). Assembled NPs were then buffer exchanged into phosphate-buffered saline (PBS) by dialysis at 4°C overnight, followed by a second dialysis step for a minimum of 4 h, using a Slide-A-Lyzer MINI dialysis device (20 kDa molecular weight cutoff; ThermoFisher Scientific). Nanoparticle concentrations were determined by the Nanodrop method using the particles peptidic molecular weight and extinction coefficient. To get these values, first the molecular weight and extinction coefficient of the SOSIP-I53-50A.1NT1 and I53-50B.4PT1 components were obtained by filling in their amino acid sequence in the online Expasy software (ProtParam tool). The peptidic mass or extinction coefficient of SOSIP-I53-50NP was then calculated by summing the obtained peptidic masses or extinction coefficient, respectively, of each component of the NP.

### SDS-PAGE and BN-PAGE analyses

For SDS-PAGE and BN-PAGE analyses, 2 µg of SOSIP trimers, or equimolar amounts of SOSIP-ferritin (2.5 µg) and SOSIP-I53-50 (3.2 µg) NPs, were run over Novex Wedge well 4–12% Tris-Glycine and NuPAGE 4–12% Bis-Tris and polyacrylamide gels (both from Invitrogen), respectively, as described earlier^29,30^. Subsequently, gels were run as per manufacturer’s protocol and then stained with PageBlue Protein Staining Solution (Thermo Scientific) or the Colloidal Blue Staining Kit (Life Technologies), respectively.

### Enzyme-linked immunosorbent assay (ELISA)

StrepTactinXT ELISA assays were performed as described previously^29,30^ with few modifications. StrepTactinXT coated microplates (IBA GmbH, Göttingen, Germany) do not require any functionalization or blocking steps prior to protein immobilization. Briefly, 100 µL of Twin-Strep-Tagged purified 329 SOSIP protein in TBS (1 µg/mL) were dispensed in the corresponding wells for protein immobilization by a 2 h incubation at room temperature. Subsequent steps to measure binding of the test antibodies were performed similarly to previously described^30^. Briefly, following a double wash step with TBS to remove unbound proteins, serial dilutions of test primary antibodies or immunized rabbit sera in Casein Blocker were added and incubated for 2 h. After 3 washes with TBS, HRP-labeled goat anti-human IgG (Jackson Immunoresearch) diluted 1:3000 in casein blocker was added and incubated for 1 h, followed by 5 washes with TBS/0.05% Tween20. Plates were developed with o-phenylenediamine substrate (Sigma-Aldrich, #P8787) in 0.05 M phosphate-citrate buffer (Sigma-Aldrich, #P4809) pH 5.0, containing 0.012% hydrogen peroxide (Thermo Fisher Scientific, #18755). Absorbance was measured at 490 nm to obtain the binding curves.

### Biolayer interferometry (BLI)

The BLI assay was performed as described earlier^29,30^, using an Octet K2 (ForteBio) device at 30°C and 1000 rpm agitation. Briefly, test antibodies diluted in kinetics buffer (PBS/0.1% bovine serum albumin/0.02% Tween20) were loaded on protein A sensors (ForteBio) to an interference pattern shift of 1 nm. Sensors were equilibrated in kinetics buffer for 60 s to obtain a baseline prior to protein association. Subsequently, purified SOSIP trimers diluted in kinetics buffer (100 nM) were allowed to associate and dissociation for 300 s. Binding data was pre-processed and exported using the Octet software.

### Nano Differential Scanning Fluorimetry (nanoDSF)

Protein thermostability was evaluated with a Prometheus NT.48 instrument (NanoTemper Technologies). Proteins at a concentration of 1 mg/mL were loaded to the grade capillaries and the intrinsic fluorescence signal was measured while temperature was increased by 1 °C/min, with an excitation power of 40%. The temperature of onset (T_onset_) and temperature of melting (T_m_) were determined using the Prometheus NT software.

### Site-specific glycan analysis using mass spectrometry

100 µg aliquots of each sample were denatured for 1h in 50 mM Tris/HCl, pH 8.0 containing 6 M of urea and 5 mM dithiothreitol (DTT). Next, Env samples were reduced and alkylated by adding 20 mM iodoacetamide (IAA) and incubated for 1h in the dark, followed by a 1h incubation with 20 mM DTT to eliminate residual IAA. The alkylated Env samples were buffer exchanged into 50 mM Tris/HCl, pH 8.0 using Vivaspin columns (10 kDa) and three of the aliquots were digested separately overnight using trypsin, chymotrypsin (Mass Spectrometry Grade, Promega) or alpha lytic protease (Sigma Aldrich) at a ratio of 1:30 (w/w). The next day, the peptides were dried and extracted using an Oasis HLB µElution Plate (Waters).

The peptides were dried again, re-suspended in 0.1% formic acid, and analyzed by nanoLC-ESI MS with an Easy-nLC 1200 (Thermo Fisher Scientific) system coupled to a Fusion mass spectrometer (Thermo Fisher Scientific) using stepped higher energy collision-induced dissociation (HCD) fragmentation. Peptides were separated using an EasySpray PepMap RSLC C18 column (75 µm × 75 cm). A trapping column (PepMap 100 C18 3μM 75μM × 2cm) was used in line with the LC prior to separation with the analytical column. The LC conditions were as follows: 275-minute linear gradient consisting of 0-32% acetonitrile in 0.1% formic acid over 240 minutes followed by 35 minutes of 80% acetonitrile in 0.1% formic acid. The flow rate was set to 300 nL/min. The spray voltage was set to 2.5 kV and the temperature of the heated capillary was set to 55 °C. The ion transfer tube temperature was set to 275 °C. The scan range was 375−1500 m/z. The stepped HCD collision energies were set to 15, 25 and 45% and the MS2 for each energy was combined. Precursor and fragment detection were performed using an Orbitrap at a resolution MS1 = 100,000. MS2 = 30,000. The AGC target for MS1 =4e5 and MS2 =5e4 and injection time: MS1 =50ms MS2 =54ms.

Glycopeptide fragmentation data were extracted from the raw file using Byos (Version 4.6; Protein Metrics Inc.). The glycopeptide fragmentation data were evaluated manually for each glycopeptide; the peptide was scored as true-positive when the correct b and y fragment ions were observed along with oxonium ions corresponding to the glycan identified. The MS data was searched using the Protein Metrics 38 insect N-glycan library. The relative amounts of each glycan at each site as well as the unoccupied proportion were determined by comparing the extracted chromatographic areas for different glycotypes with an identical peptide sequence. All charge states for a single glycopeptide were summed. The precursor mass tolerance was set at 4 ppm and 10 ppm for fragments. A 1% false discovery rate (FDR) was applied. The relative amounts of each glycan at each site as well as the unoccupied proportion were determined by comparing the extracted ion chromatographic areas for different glycopeptides with an identical peptide sequence. Glycans were categorized according to the composition detected.

Glycans were categorized according to the composition detected. HexNAc(2)Hex(9−4) was classified as M9 to M4. Any of these compositions containing fucose were classified as fucosylated mannose (FM). HexNAc(3)Hex(5−6)X was classified as Hybrid with HexNAc(3)Fuc(1)X classified as Fhybrid. Complex-type glycans were classified according to the number of processed antenna and fucosylation. Complex-type glycans were categorized according to the number of N-acetylhexosamine monosaccharides detected, that do not fit in the previously defined categories. If all of the compositions have a fucose they are assigned into the (F) categories. As this fragmentation method does not provide linkage information compositional isomers are group, so for example a triantennary glycan contains HexNAc 5 but so does a biantennary glycans with a bisect. Any glycan containing at least one sialic acid was counted as sialylated.

### Negative-stain electron microscopy (nsEM)

Purified 329 SOSIP Env or SOSIP ferritin NP or SOSIP-I53-50 NP proteins were diluted to 0.03 - 0.05 mg/ml in PBS before grid preparation. A 3 µL drop of diluted protein (∼0.025 mg/ml) was applied to previously glow-discharged, carbon-coated grids for ∼60 s, blotted and washed twice with water, stained with 0.75 % uranyl formate, blotted, and air-dried. Between 30 and 50 images were collected on a Talos L120C microscope (Thermo Fisher) at 73,000 magnification and 1.97 Å pixel size. Relion-3.1^43^ or Cryosparc v4.5.1^44^ were used for particle picking and 2D classification.

### CryoEM sample preparation, data acquisition and data analysis

Three µL of Purified SOSIP I53-50 NP sample at the concentration of 0.5 mg/ml was applied onto a freshly glow-discharged (PLECO easiGLOW) 300 mesh, 1.2/1.3 C-Flat grid (Electron Microscopy Sciences). After 20 s of incubation, grids were blotted for 3 s at 0 blot force and vitrified using a Vitrobot IV (Thermo Fisher Scientific) under 22°C with 100% humidity. Single-particle Cryo-EM data was collected on a 200 kV Talos Arctica transmission electron microscope (ThermoFisher Scientific) equipped with Gatan K3 direct electron detector behind a 20 eV slit width energy filter. Multi-frame movies were collected at a pixel size of 1.1 Å per pixel with a total dose of 58.3 e/Å^2^ at defocus range of −0.5 to −2.4 µm. ∼3142 cryoEM movies were motion-corrected by Patch motion correction implemented in Cryosparc v4.5.1^44^. Motion-corrected micrographs were corrected for contrast transfer function using Cryosparc’s implementation of Patch CTF estimation. Micrographs with poor CTF fits were discarded using CTF fit resolution cutoff of ∼6.0 Å. Particles were picked using a Blob picker, extracted, and subjected to an iterative round of 2D classification. Particles belonging to the best 2D classes with secondary structure features were selected for two classes of Ab-initio reconstruction. Particles belonging to the best Ab-Initio class were refined in non-uniform 3D refinement with per particle CTF and higher-order aberration correction turned on and applying Icosahedral (I1) symmetry to generate cyoEM density map. Model for I53-50A and I53-50B nanoparticle (PDB:7SGE) was docked into the map using Chimera1.7.1 ^45^ fit in map function.

### Rabbit immunizations

The rabbit immunization was outsourced to a contract research organization (CRO) named Liveon Biolabs Private Limited, Bengaluru, Karnataka, India. The immunization studies described here were carried out on female naive New Zealand White rabbits of 2.0–2.5 kg and age 4 months. The use of animals for this study was approved by Liveon Biolabs Private Limited IAEC. IAEC approved Protocol No.: LBPL-IAEC-008-01/2021 with study number: LBPL/NG-1736 (EF). All immunization procedures complied with animal ethical regulations and protocols of the Liveon Biolabs Private Limited IAEC committee. For all immunogens (329 SOSIP, SOSIP-ferritin NP and SOSIP-I53-50 NP), groups of four rabbits were given two intramuscular immunizations in each quadriceps at weeks 0, 4, and 20. The immunization mixture involved 20 µg of SOSIP trimers, or an equimolar amount presented as SOSIP-ferritin NPs (25 µg) or as SOSIP-I53-50 NPs (32 µg), formulated in AddaVax adjuvant (1:1 v/v). Dose calculations were based on the peptidic molecular weight of the proteins (thus disregarding glycans), which were obtained essentially as described ^26,27^. The rabbits were bled at weeks 0, 4, 6, 16, 20 and 22.

### HIV-1 pseudovirus generation

The HIV-1 pseudoviruses were produced in HEK293T cells as described earlier^17,46–48^, by co-transfecting the corresponding full HIV-1 gp160 envelope plasmid and a pSG3ΔEnv backbone plasmid. Briefly, 1×10^5^ cells in 2 mL complete DMEM (10 % fetal bovine serum (FBS) and 1 % penicillin and streptomycin antibiotics) were seeded per well of a 6 well cell culture plate (Costar) the day prior to transfection. For transfection, envelope (1.25 μg) and delta envelope plasmids (2.50 μg) were mixed in a 1:2 ratio in Opti-MEM (Gibco), with a final volume of 200μl per well, and incubated for 5 minutes at room temperature. Next, 3 μl of PEI-Max transfection reagent (Polysciences) (1 mg/ml) was added to this mixture prior to further incubation for 15 min at room temperature. This mixture was then added dropwise to the HEK 293T cells supplemented with fresh complete DMEM growth media and incubated at 37 °C for 48 h. Pseudoviruses were then harvested by filtering cell supernatants with 0.45 μm sterile filters (mdi), aliquoted and stored at −80 °C until usage.

### HIV-1 neutralization assays

The neutralizing activity rabbit immune sera was tested against autologous and heterologous pseudoviruses, by performing neutralization assays as described earlier^49–51^. Neutralization was measured as a reduction in luciferase gene expression after a single round of infection of TZM-bl cells (NIH AIDS Reagent Program) with HIV-1 Env pseudoviruses. The TCID_50_ of the HIV-1 pseudoviruses was calculated and 200 TCID_50_ of the virus was used in neutralization assays by incubating with 1:3 serially diluted rabbit sera starting at 1:20 dilution. After that, freshly trypsinized TZM-bl cells in growth medium (complete DMEM with 10% FBS and 1% penicillin and streptomycin antibiotics) containing 50 μg/ml DEAE Dextran at 10^5^ cells/well were added and plates were incubated at 37°C for 48 h. Virus controls (cells with HIV-1 virus only) and cell controls (cells without virus and antibody) were included. MuLV was used as a negative control. After the incubation of the plates for 48 h, luciferase activity was measured using the Bright-Glow Luciferase Assay System (Promega). ID_50_ for antibodies were calculated from a dose-response curve fit with a non-linear function using the GraphPad Prism 9 software (San Diego, CA). All neutralization assays were repeated at least 2 times, and data shown are from representative experiments.

### Statistical analysis

Graphpad Prism version 9.0 was used for all statistical analyses.

## DATA AVAILABILITY

All data generated or analysed during this study are included in this article and its supplementary information files. The data generated and analysed during the current study available from the corresponding author on reasonable request.

## ACKNOWLEDGEMENTS

This study was funded by Department of Biotechnology, India (BT/PR30120/MED/29/1339/2018) grant awarded to K.L. This study was supported by EMBO through the Short-Term Fellowship (STS-7814) awarded to S.K. This project has received funding from the European Union’s Horizon 2020 research and innovation program under grant agreement No. 681137 (to R.W.S. and M.C.). This work was also supported by the U.S. National Institutes of Health Grant P01 AI110657 (to R.W.S.); by the International AIDS Vaccine Initiative (IAVI); by the Bill and Melinda Gates Foundation through the Collaboration for AIDS Vaccine Discovery (CAVD), grants OPP1111923 and OPP1132237 (to R.W.S.); by the Aids Funds Netherlands, Grant #2016019 (to R.W.S.); and by the Fondation Dormeur, Vaduz (to R.W.S.). R.W.S. is a recipient of a Vici grant from the Netherlands Organization for Scientific Research (NWO). The funders had no role in study design, data collection and analysis, decision to publish, or preparation of the manuscript. We are very much thankful to NIH AIDS reagent program for HIV-1 research reagents, Neutralizing antibody consortium (NAC), IAVI, USA for HIV-1 neutralizing antibodies donated by Michel Nussenzweig, Hermann Katinger, Mark Connors, James Robinson, Dennis Burton, John Mascola, Peter Kwong, and William Olson. The nsEM and cryo-EM datasets on Talos Arctica were collected at Robert P. Apkarian Integrated Electron Microscopy Core (IEMC) at Emory University, Atlanta. We thank the IEMC staff members for support in data collection.

## AUTHOR CONTRIBUTIONS

S.K., I.d.M-S., R.W.S. and K.L. conceived and designed experiments. S.K., I.d.M-S., S.S., M.L.N., J.D.A., T.P.L.B., Y.V., L.J., and A.P. performed the experiments. S.K., I.d.M-S., S.S., J.D.A., E.A.O., A.P., M.C., R.W.S. and K.L. analyzed and interpreted data. S.K. and S.S. organized the rabbit immunization studies. S.K., I.d.M-S., R.W.S. and K.L. wrote the manuscript. All authors reviewed, edited and/or provided input to the manuscript.

